# Rabbit Hindlimb Kinematics and Ground Contact Pressure During the Stance Phase of Hopping Gait

**DOI:** 10.1101/2022.02.02.478798

**Authors:** Patrick T. Hall, Caleb Stubbs, David E. Anderson, Cheryl B. Greenacre, Dustin L. Crouch

**Author notes:** Corresponding Author: Patrick Hall.

## Abstract

Though the rabbit is a common animal model in musculoskeletal research, there is very limited data reported on healthy rabbit biomechanics. Our objective was to quantify the normative hindlimb biomechanics of six New Zealand White rabbits (3 male, 3 female) during the stance phase of hopping gait. We measured biomechanics by synchronously recording sagittal plane motion and ground contact pressure using a video camera and pressure mat, respectively. Both foot angle (i.e., angle between foot and ground) and ankle angle curves were unimodal. The peak ankle dorsiflexion angle was 65.9°±12.7° and occurred at 39% stance, while the peak ankle plantarflexion angle was 136.9°±7.8° at toe-off. Minimum and maximum foot angles were 16.6°±6.4° at 12% stance and 125.4°±4.0° at toe-off, respectively. During stance, the knee joint center gradually progressed 4.7 cm downward and 18.1 cm forward, on average. The maximum vertical ground reaction force and contact area, both averaged across rabbits, were 42.5 ± 11.4 %BW and 7.5 ± 1.8 cm^2^, respectively. Our study confirmed that rabbits exhibit a plantigrade gait pattern, similar to humans. Future studies can reference our data to quantify the extent to which orthopedic interventions affect rabbit biomechanics.

## Introduction

The rabbit is a common animal model in musculoskeletal research to, for example, test new potential clinical interventions or study the response of tissues to mechanical stimuli. Some interventions would be expected to affect the motor function of the hindlimb ankle and foot. Examples of such interventions include tenotomy of ankle dorsiflexor muscles (Abrams et al. 2000) or the Achilles tendon (Nagasawa et al. 2008; Reddy et al. 1998), immobilization of the knee and/or ankle joint (Gossman et al. 1986; Ponten & Friden 2008; Sjostrom et al. 1979), and release of tendon retinacula to manipulate muscle-tendon moment arms (Koh & Herzog 1998b; Reddy & Gupta 2007). Recently, we adopted a rabbit model to test the feasibility of a new type of ankle-foot prosthesis (Hall et al. 2021). For these interventions, quantifying their effect on motor function will be valuable, if not essential, for achieving clinical translation.

One way to quantify motor function is by measuring biomechanical variables, such as kinematics and ground reaction forces. The effect of an intervention could be determined by comparing biomechanical data between animals that did and did not receive the intervention. Unfortunately, the hindlimb biomechanics of healthy rabbits has not been well characterized. We are aware of only one previous study in which multi-sample data were limited to knee and ankle joint moments during hopping gait (Gushue et al. 2005). A more comprehensive dataset on normative rabbit hindlimb biomechanics in the literature would (1) make it easier for researchers to determine the effects of experimental interventions on motor function, (2) reduce the number of healthy animals used as experimental controls, and (3) increase basic understanding about animal locomotion.

Two common measures of hindlimb biomechanics that have not been reported for a multi-sample rabbit cohort are joint kinematics and ground contact pressure. These variables, in some respects, are similar between the rabbit hindlimb and human lower extremity. For example, the rabbit hindlimb (Bensley 1910; Kimura 1996), like the human lower extremity (Grasso et al. 1998), is considered to exhibit a plantigrade kinematic gait pattern in which most or all of the plantar surface of the foot contacts the ground during the stance phase. This differs from other common animal models that exhibit either a digitigrade (e.g., cats, dogs) (Barbeau & Rossignol 1987) or unguligrade (e.g. pigs and goats) (Polly 2007) gait pattern. Likewise, the net vertical ground reaction force (i.e., the integral of ground contact pressure across the contact area) during stance phase has a bimodal pattern in both rabbits (Gushue et al. 2005), and humans (Wannop et al. 2012). Thus, in interventional studies, joint kinematics and ground contact pressure results from rabbits may be more translateable to humans than those from other common animal models.

The goal of our study was to quantify hindlimb ankle and foot kinematics and ground contact pressures during the stance phase of hopping gait in healthy rabbits (i.e., rabbits that have not received an experimental intervention). Summary results are presented below, and the raw data are available online as supplementary data. The new biomechanics data will serve as a valuable reference for interventional studies involving rabbits and improve our basic understanding about rabbit locomotion.

## Materials & Methods

All animal procedures were approved by the University of Tennessee, Knoxville Institutional Animal Care and Use Committee (protocol #2637). We used a cross-sectional study design, measuring biomechanics in one test session from both hindlimbs of six (3 male, 3 female) healthy, standard laboratory New Zealand White (NZW) rabbits (Charles River Laboratories, Wilmington, MA). At the time of testing, the rabbits were 16 weeks old and weighed 2.7 ± 0.33 kg. Each rabbit was one experimental unit. Since this was not a clinical trial, there were no separate groups (i.e., control), no inclusion criteria, and no power analysis to compute sample size. Rabbits were housed individually in adjacent crates, fed ad libitum with a standard laboratory diet and Timothy hay, and given daily enrichment and positive human interaction. After this study, the rabbits were subsequently used in another study to test a novel orthopedic implant (Hall et al. 2021).

The setup for our locomotor measurement system included several components (Fig. 1, 1-3). A pressure mat (Very High Resolution Walkway 4, Tekscan, Inc., South Boston, MA) was used to record pressure data at 60 Hz. We taped 320-grit sandpaper to the top of the smooth pressure-sensing area to prevent the rabbits from slipping. We placed a 3-kg weight on the sandpaper-covered mat and calibrated the mat through the Tekscan software calibration program. The mat was placed inside a clear acrylic tunnel to guide the rabbits across the pressure mat. The mat width (11.2 cm) permitted only unilateral pressure measurements; therefore, to record data for both hindlimbs, we laterally offset the mat in the tunnel and had the rabbit hop across the mat in both directions, as described below. A camera (1080P HD Webcam, SVPRO), placed three feet away from the clear acrylic panel, captured video at 60 Hz. Video and pressure mat data were synchronized using the Tekscan Walkway software (Tekscan, South Boston, MA).

**Figure 1.**
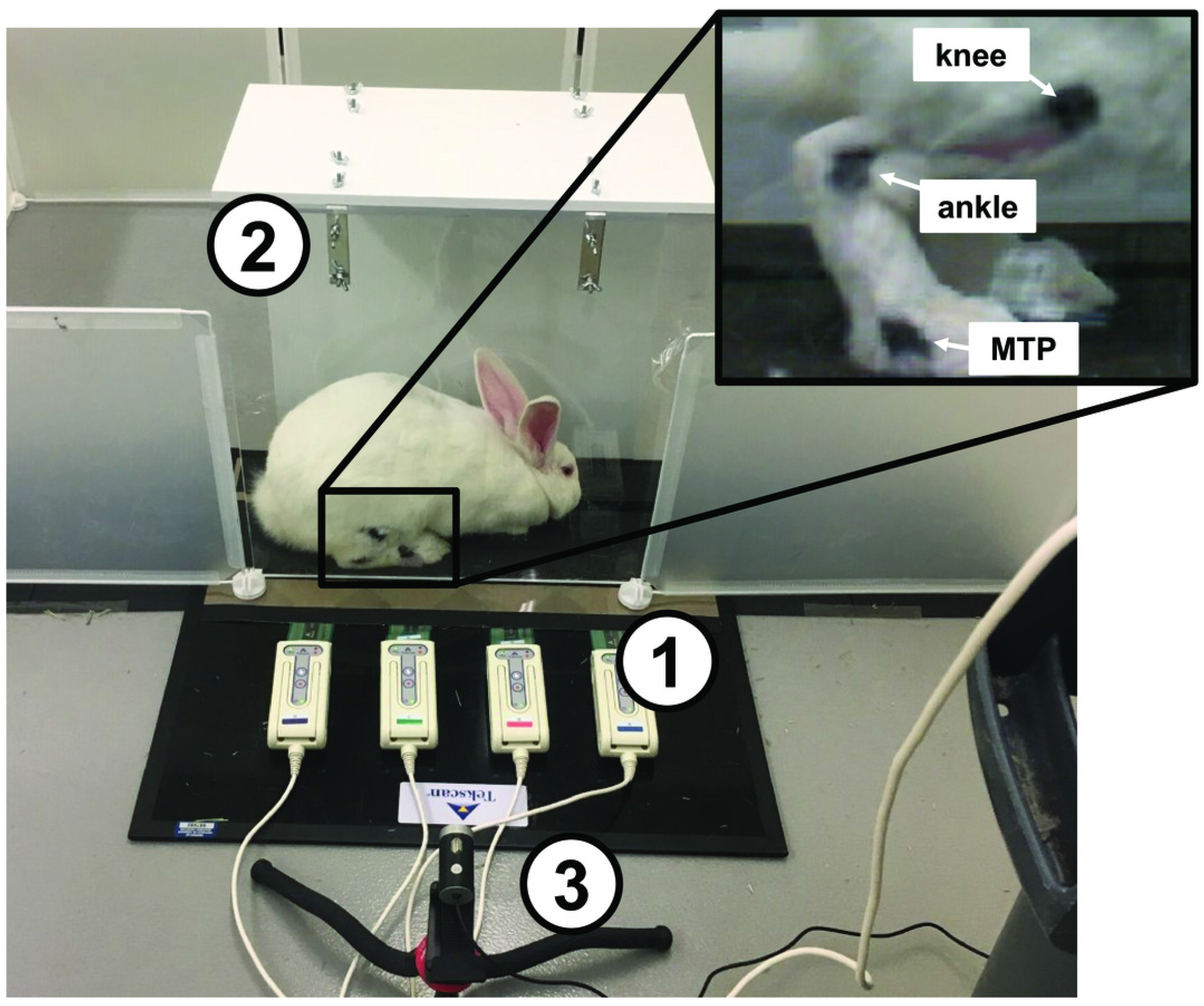
Testing setup used to collect motion capture data. **(1)** Tekscan Very HR Walkway 4 for measuring pressure data. **(2)** Acrylic tunnel for guiding rabbits across pressure mat. **(3)** 60 Hz Camera for recording sagittal plane kinematics. **(4)** Black ink marks at approximate joint centers based on bony landmarks.

During a two-week acclimation period prior to testing, rabbits were trained to hop through the acrylic tunnel when given negative reinforcement (i.e., prodding). Then, at the beginning of the test session, we shaved both hindlimbs and marked the metatarsophalangeal (MTP), ankle, and knee joint centers on the lateral side of each hindlimb with black ink (Fig. 1, 4) to facilitate calculation of joint kinematics from the videos. After marking, we placed the rabbit into a pen with the acrylic tunnel and pressure mat. Each trial began when the rabbit entered the tunnel. A trial was deemed successful if the rabbit continued the hopping motion through the entire length of the tunnel without stopping. The rabbits completed 5 trials of hopping through the tunnel in each direction while we synchronously recorded pressure and video data for each trial. The direction of hopping is a potential confounder, which we address as a limitation in the Discussion section.

Data from all rabbits were included in the analysis. We calculated joint angles from the video frames corresponding to stance phase using a custom script in MATLAB (Mathworks, Natick, MA). The program used a bottom-hat morphology filter to distinguish the black ink marks and calculate the centroid of each marker position (Fig. 2). Then, frame-by-frame, we visually verified the marker centroids and, if the centroid location appeared inaccurate, corrected the location by manually approximating the centroid from the still frame. This method assumed that the sagittal plane orientation of the foot and shank was aligned with the line segments connecting the centroids; this was a reasonable assumption given the arrangement of the camera with respect to the acrylic tunnel. Finally, we calculated the angle between the foot segment and ground (i.e., foot angle) and between the foot and shank segments (i.e., ankle angle) throughout stance (Fig. 2). Since the camera was approximately level with the ground, we defined the ground as the horizontal line that intersected the MTP joint. Ankle angles >90° and <90° corresponded to plantarflexion and dorsiflexion, respectively.

**Figure 2.**
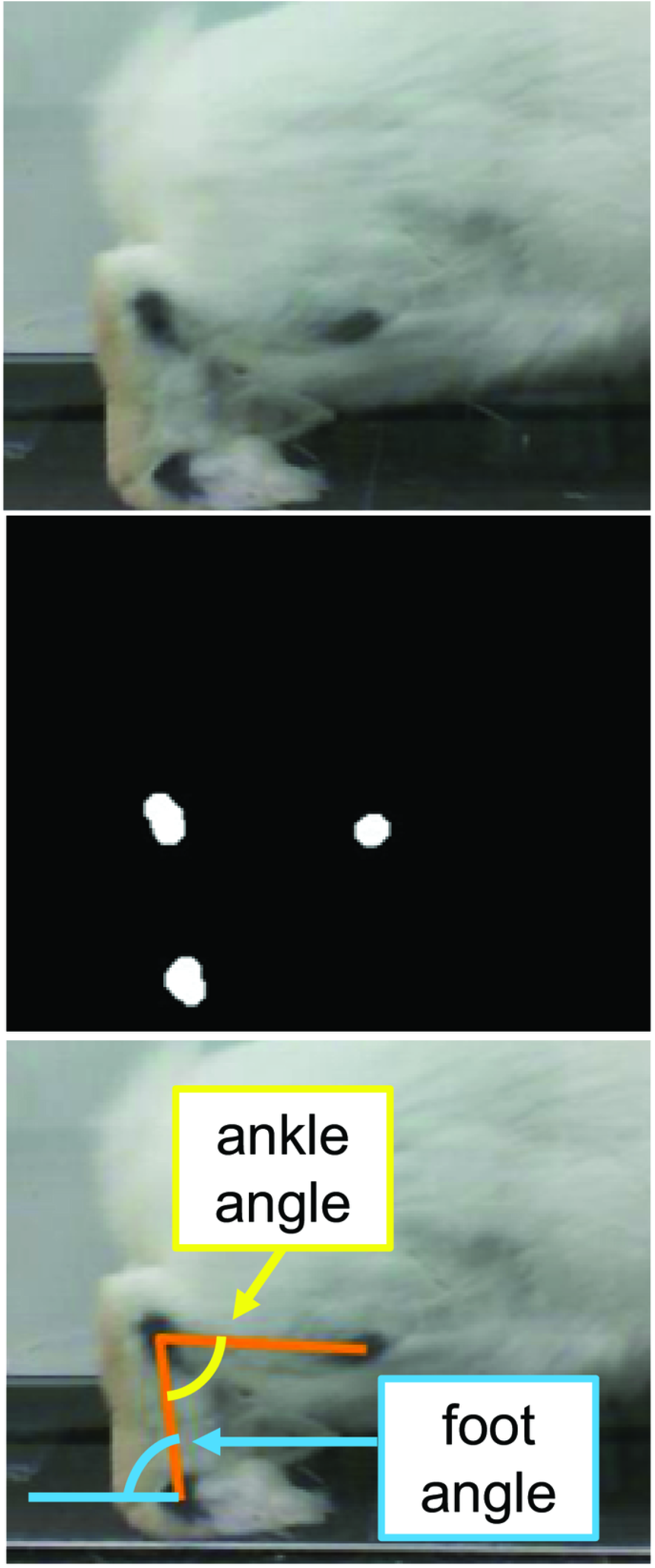
Motion capture analysis from sagittal plane videos for calculating hindlimb kinematics. A frame-by-frame top-hat morphology permitted detection of joint markers. Vectors representing limb segments (orange lines) were drawn between the centroids of the joint markers. Using the dot product and the segment vectors, we calculated the ankle angle (yellow) and foot angle (i.e., the angle between the foot and ground) (blue). We considered the ground (dashed blue) to be a horizontal line that intersected the MTP joint.

Evaluating joint angles separately for each joint provides an incomplete picture of rabbit hindlimb kinematics. This is because the hindlimb is a kinematic chain, so the positions and orientations of proximal limb segments depend on those of distal limb segments. Therefore, we calculated the knee joint center position in the sagittal plane from the foot and ankle angles. To permit comparison among rabbits of different sizes, we applied the joint angles to a generic planar kinematic model of the rabbit hindlimb (Fig. 3, A). The foot (ankle to MTP joint) and shank (knee to ankle) segments of the model were 6.7 cm and 10.0 cm, respectively, to approximate the anatomical lengths of the two segments in our rabbits. For each rabbit, we computed the time-series “trajectory” of knee joint position throughout stance for each trial.

**Figure 3.**
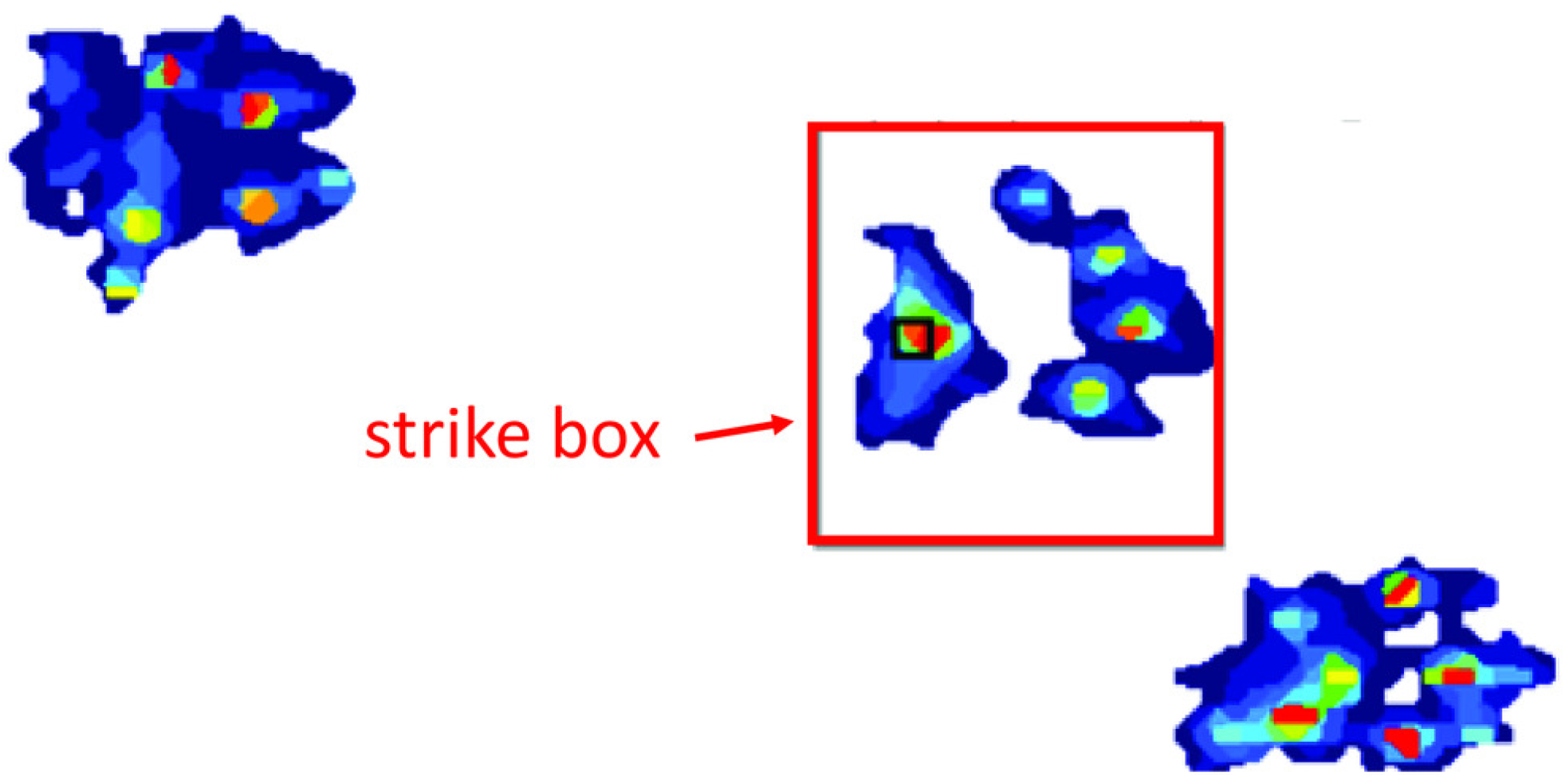
Example of strike box used to isolate a single foot from ground contact pressure data using Tekscan Walkway software.

Ground contact pressure data were processed using the Tekscan Walkway software. Each hind foot was isolated by drawing a strike box around the foot contact location, as indicated by the pressure data (Fig. 4). From the pressure data in the strike box and for each time point, we computed (1) average pressure, (2) contact area, and (3) net vertical ground reaction force (vGRF). The contact area was calculated as the total geometric area within the strike box for which the pressure was greater than zero. The vGRF was calculated as the product of foot contact area and the average pressure across the contact area; vGRF data were expressed as a percentage of body weight (%BW). We also computed the total contact area over the entire stance phase. The duration of the stance phase for each limb was computed as the length of time for which pressure was greater than zero within the limb's strike box.

**Figure 4.**
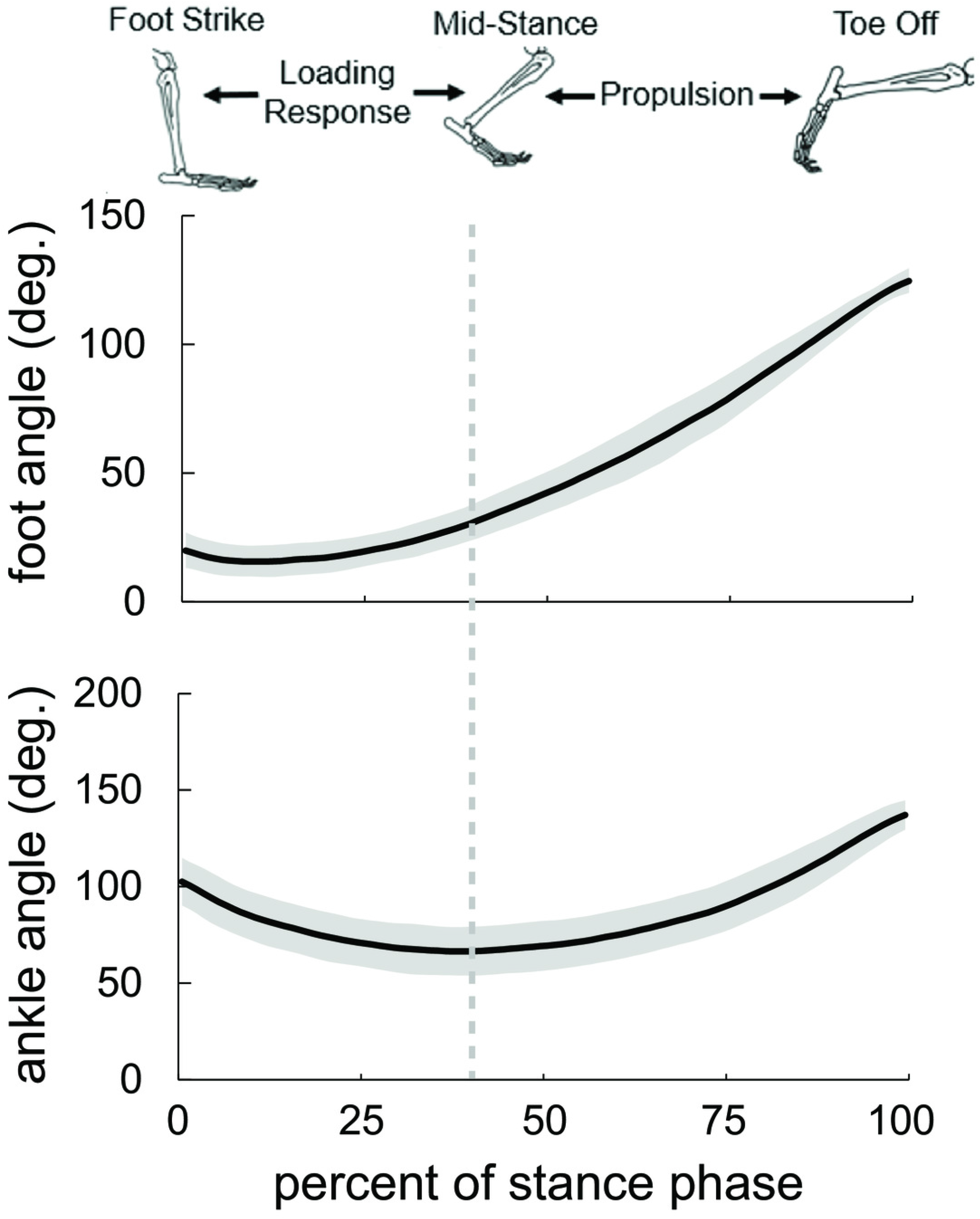
Joint kinematics of the rabbit hindlimb during stance phase. Mean and ±1 standard deviation represented by black line and grey shaded region, respectively. Both foot angle (top) and ankle angle (bottom) exhibited a unimodal curve. Consistent with human gait, we divided rabbit stance into “loading response” and “propulsion” sub-phases at the maximum ankle dorsiflexion (i.e., smallest value) angle.

In each trial, the data were calculated separately for each side (i.e., right and left). Since the sides were measured independently, each side was considered as an independent sample in calculations of the mean and standard deviation (i.e., n=12).

## Results

At foot strike, the ankle angle exhibited a similar pattern to that of humans (Li & Hsiao-Wecksler 2013), with the ankle becoming increasingly dorsiflexed for the first 39% of stance, commonly referred to as “loading response”; for the remaining 61% of stance, considered the “forward propulsion” sub-phase, the ankle became increasingly plantarflexed (Fig. 4). The peak ankle dorsiflexion angle was 65.9°±12.7°, while the ankle was plantarflexed at both foot strike (101.2°±13.7°) and toe-off (136.9°±7.8°). The foot maintained a positive angle throughout stance, starting at 21.1°±7.3°, then decreasing to a minimum foot ankle of 16.6°±6.4° at 12% stance. Thereafter, the foot ankle gradually increased up to 125.4°±4.0° at toe-off, indicating that the foot rotated beyond a vertical orientation (90°). The standard deviation of the foot and ankle angles, averaged over the stance phase, was 7.7° and 12.7° degrees, respectively.

Despite both foot and ankle angles exhibiting a unimodal curve, the knee joint center trajectory was nearly linear (Fig. 5). During stance, the knee joint center gradually progressed 4.7 cm downward and 18.1 cm forward, on average. The standard deviation of the knee joint center position, averaged across the stance phase, was 2.1 cm.

**Figure 5.**
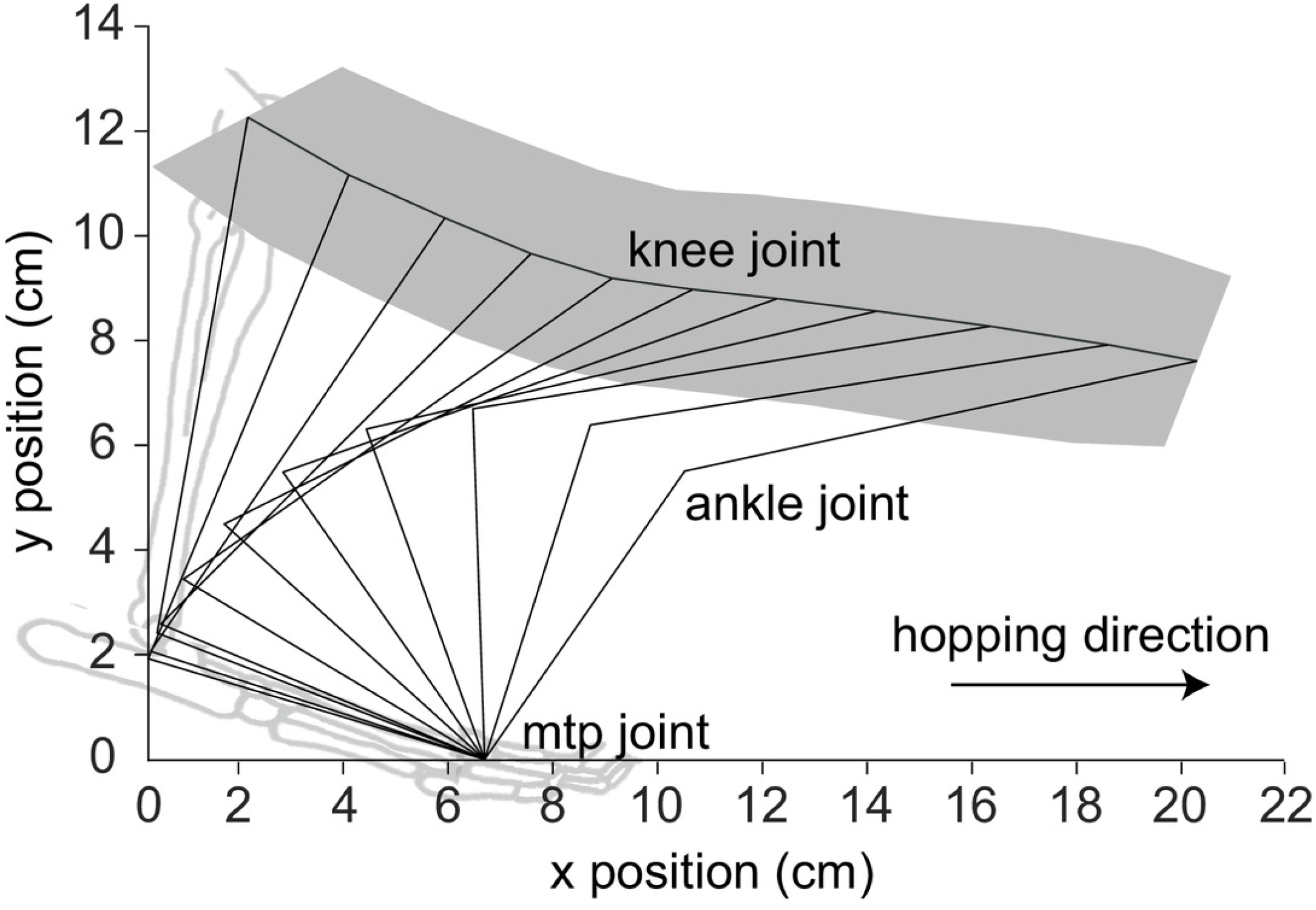
Knee joint center position during stance phase. Mean and ±1 standard deviation represented by black line and grey shaded region, respectively. The knee joint gradually progressed forward and downward due to the combination of foot and ankle joint angles.

The maximum vGRF was 42.5±11.4 %BW, and the maximum contact area was 7.5±1.8 cm^2^ (Fig. 6). Both vGRF and contact area exhibited similar bimodal trends during stance, with the first peak occurring at approximately 25% of stance and the second occurring at approximately 75% of stance. In fact, vGRF and contact area were strongly correlated, with a Pearson's correlation coefficient of 0.986. Because of the correlation between vGRF and contact area, the foot experienced a nearly constant average pressure of about 0.15 kg/cm^2^ through the middle 80% of stance.

**Figure 6.**
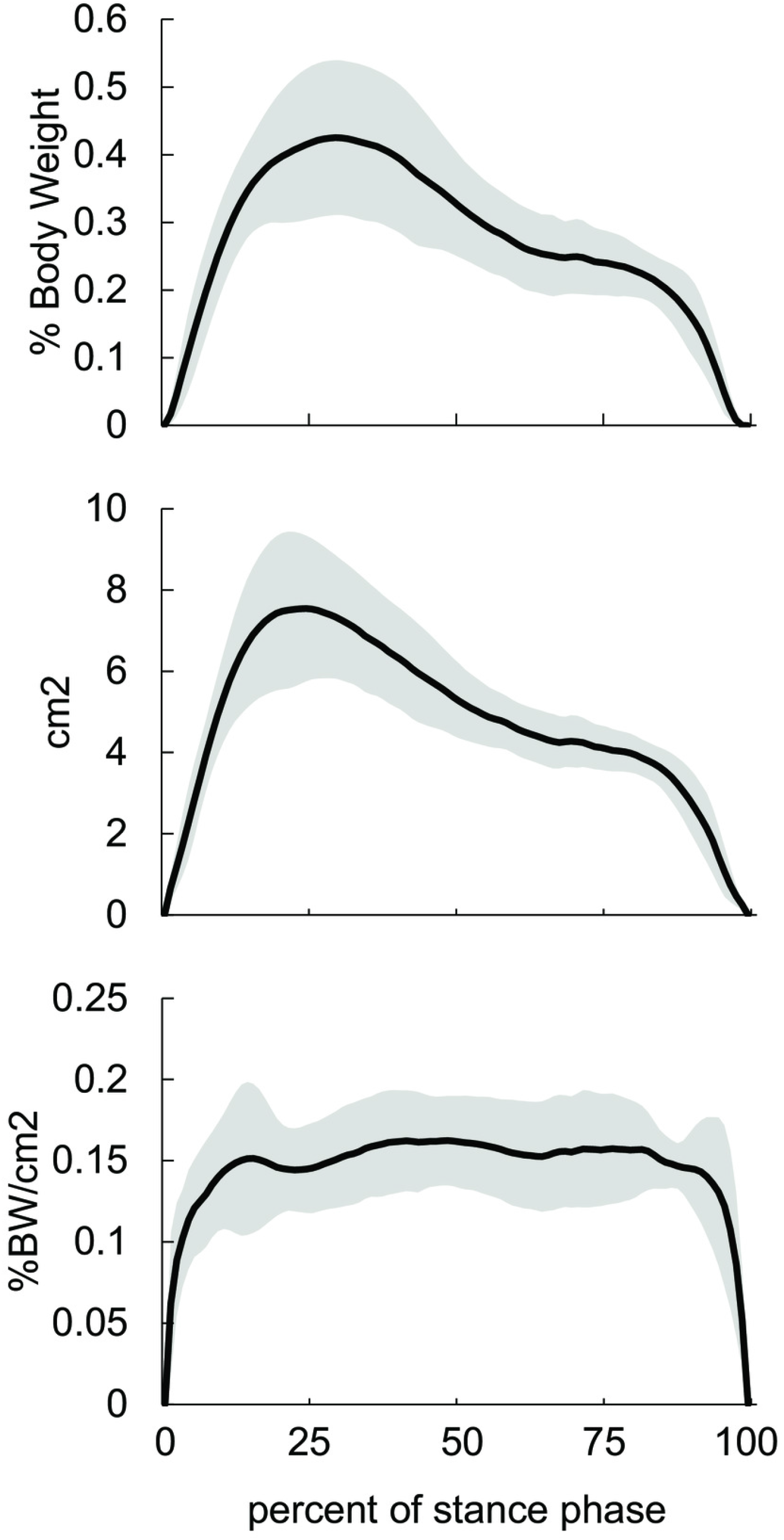
Characteristics of ground contact of one hindlimb during stance phase. Mean and ±1 standard deviation represented by black line and grey shaded region, respectively. **(Top)** Vertical ground reaction force (vGRF) expressed as a percentage of body weight. vGRF had a bimodal pattern, with peaks at approximately 25% and 75% of stance. **(Middle)** Ground contact area, computed as the total surface area for which measured pressure was greater than zero. Similar to vGRF, contact area had a bimodal pattern. vGRF and contact area were strongly correlated (Pearson's r= 0.986). **(Bottom)** Average ground contact pressure. Given the correlation between vGRF and contact area, pressure was nearly constant over the middle 80% of stance.

The spatial distribution of ground contact pressure indicated that, relative to the foot, the contact area progressed from the caudal to cranial aspect of the plantar surface during stance (Fig. 7). At foot strike, the heel and midfoot regions of the plantar surface were the first to contact the ground. By about 25% stance, the contact area in these regions increased and expanded toward the cranial aspect (i.e., forefoot) of the plantar surface. By about 65% of stance, the contact area shifted entirely to the forefoot region.

**Figure 7.**
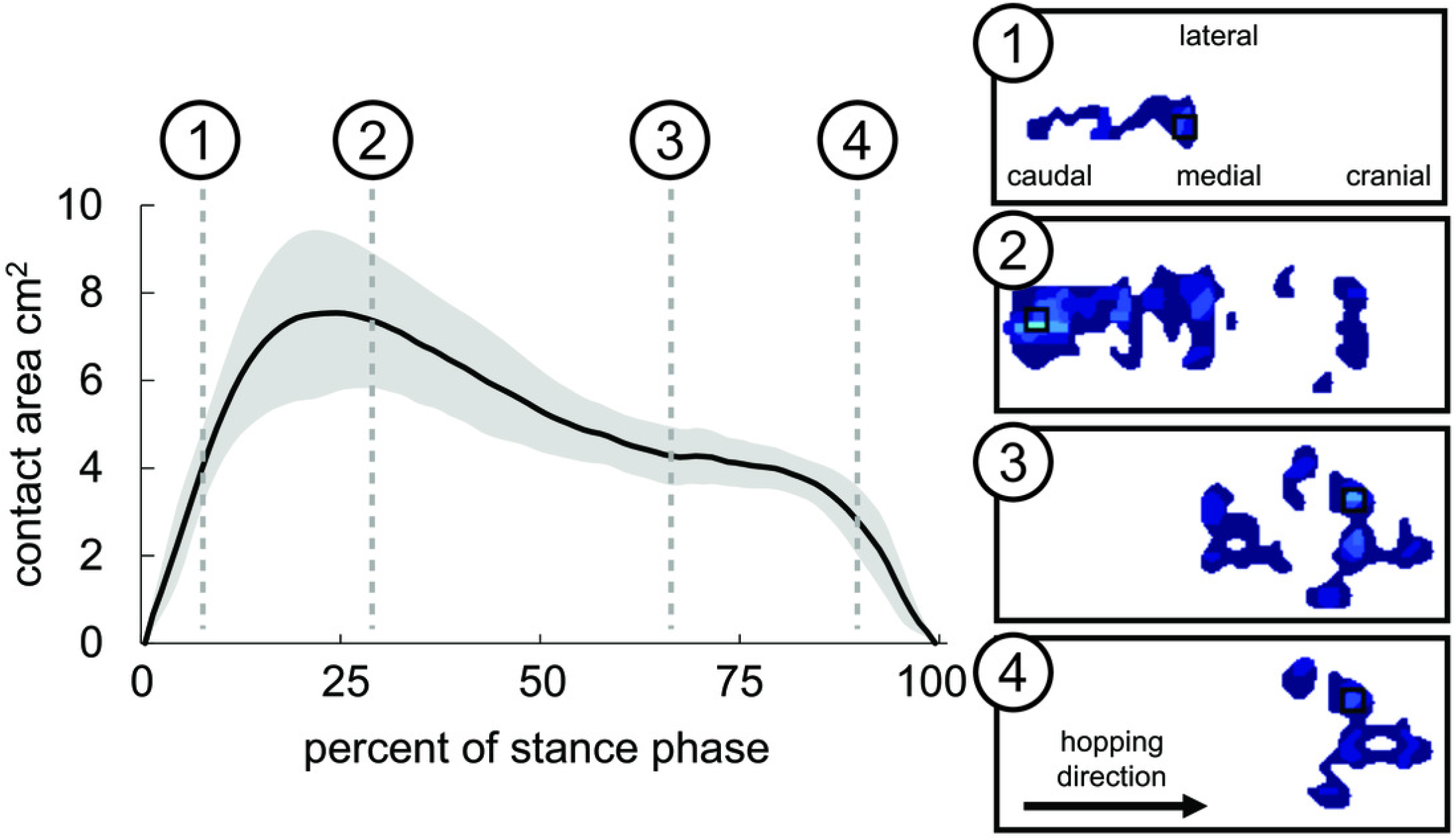
Ground contact area (left) and pressure distribution from a representative trial (right) during stance phase. Mean and ±1 standard deviation represented by black line and grey shaded region, respectively. Ground contact pressure generally progressed from the caudal to cranial direction of the foot during stance.

The stance phase duration, averaged across all trials and sides, was 0.50±0.27 seconds but had a right-skewed distribution; the median duration was 0.40 seconds.

## Discussion

Our kinematic and pressure data confirmed that rabbits exhibit a plantigrade gait pattern, loading the heel first (Fig. 5). Humans also have a plantigrade walking pattern and land with their heel first (Grasso et al. 1998). However, there are differences in kinematics between rabbits and humans. For example, foot angle during stance is more negative during human walking (Grasso et al. 1998) than rabbit hopping (Fig. 5). Such differences must be considered when attempting to translate results from animal models to humans.

The time-series vGRF data exhibited a bimodal pattern that is consistent with vGRF data reported for one rabbit (Gushue et al. 2005) and for humans during walking (Wannop et al. 2012). We expect that, as with humans, the maximum vGRF that occurs at approximately 25% of stance corresponds to the loading response triggered by the need to support body weight during landing (Winiarski & Rutkowska-Kucharska 2009). As in humans (Grasso et al. 1998), the second mode at approximately 75% of stance presumably corresponds to forward propulsion, though we could not confirm this since the pressure mat cannot record horizontal ground reaction forces.

Contact area was strongly correlated with vGRF magnitude, with a correlation coefficient of 0.986 across all trials. The correlation explains why the average pressure was nearly constant throughout stance phase. Maintaining a constant ground-paw pressure may be a locomotor strategy employed by the rabbit's sensorimotor system. However, as seen in our prior study (Hall et al. 2021), orthopedic interventions can alter the pressure between the hindlimb and ground.

We computed the sagittal-plane position of the knee joint center, which gives an indication of how the joint is moving in the global coordinate system. In contrast, joint angles represent the orientation of one local coordinate frame to another, where both frames are aligned with a limb segment. Thus, joint center position data provide important context to joint angle data by showing how local joint-level changes may affect global limb or body position. Joint center positions may even better reflect task goals than joint angles. For example, a previous study found that rats with peripheral nerve injury recovered joint position kinematics even as individual joint angles remained altered (Chang et al. 2018).

We chose an approach to measure rabbit hindlimb biomechanics that is practical for longitudinal interventional studies in rabbits. For example, we used a markerless video-based motion capture method to quantify hindlimb kinematics. This differs from the state-of-the-art method that involves tracking reflective markers using multiple infrared cameras. Reflective markers are typically placed on a subject's skin using adhesive tape, but this method is unreliable in rabbits, in our experience. This possibly explains why the previous rabbit biomechanics study (Gushue et al. 2005) attached reflective markers using transdermal bone pins which, though reliable, are relatively difficult to implement and may interfere with movement either mechanically or by causing discomfort. Bi-plane fluoroscopy can acquire detailed kinematic data (Koh & Herzog 1998a; Tinga et al. 2018) but poses a radiation safety risk to the animals and researchers in longitudinal studies. Finally, both infrared- and fluoroscopy-based methods require relatively expensive equipment that must remain stationary during testing and, thus, are not as accessible, portable, or mobile as video-based methods.

Pressure mats are also more practical for measuring ground contact forces in quadrupedal animal models than the alternative, force plates. Force plates measure the *resultant* forces along 6 degrees of freedom, which requires that only one limb contacts the plate at a time to distinguish forces among limbs (Gushue et al. 2005; Jarrell et al. 2018). Pressure mats, which measure vertical pressure and contact area, are more convenient to use with quadrupedal animals since individual limbs can be distinguished even if multiple limbs contact the mat simultaneously (de Carvalho et al. 2009; Sheldon et al. 2019; Steiner et al. 2019).

Our study was limited in several ways. First, because our pressure mat was relatively narrow (11.2 cm), we only measured kinematic and pressure data for one hindlimb at a time. In future studies we plan to use a wider pressure mat and two cameras (one on each side) to capture biomechanics data from both hindlimbs simultaneously; this data will allow us to quantify temporal relationships between sides. Second, the sample frequency (60 Hz) of our motion capture setup allowed us to capture stance phase only; in future studies we will use cameras with a higher sample frequency to also capture the high-frequency kinematics of the swing phase. Third, the kinematics we computed with our custom MATLAB script were limited to the sagittal plane. The more-advanced open-source software DeepLabCut (Mathis et al. 2018a; Mathis et al. 2018b) uses machine learning to track anatomical features automatically and calculate 3-D kinematics from markerless motion capture videos; we plan to use such software in future studies. Fourth, we tracked the motion of marked points on the skin, which are more convenient than the bone pins used in the previous rabbit biomechanics study (Gushue et al. 2005) but can move relative to underlying bony landmarks.

## Conclusions

In conclusion, we have reported select hindlimb kinematics and ground contact pressures from healthy New Zealand white rabbits. Our results showed that rabbits exhibit a plantigrade gait pattern as described in previous literature. Hindlimb ankle kinematics during stance were unimodal, similar to humans. Vertical ground reaction force (vGRF) and contact area were highly correlated, resulting in relatively constant pressure during stance. Our results add substantially to the limited existing data on rabbit hindlimb biomechanics and can be used as an experimental control to quantify the effect experimental interventions. Additionally, knowledge of rabbit hindlimb biomechanics can inform the design and development of orthopedic interventions to improve functional outcomes of patients.

## Supporting information

Kinematics and ground contact pressure data

## Acknowledgments

The authors thank Dr. Bryce Burton, Dr. Kelsey Finnie, Dr. Lori Cole, and Chris Carter for veterinary care provided for the rabbits in this study, and the Office of Laboratory Animal Care and Animal Housing Facility staffs at the University of Tennessee, Knoxville for animal care assistance. Thanks to Dr. Katrina Easton for providing constructive feedback on the manuscript. Research reported in this publication was supported by (1) the Eunice Kennedy Shiver National Institute of Child Health & Human Development of the National Institutes of Health under Award Number K12HD073945, (2) NSF CAREER Award #1944001, (3) a seed grant from the University of Tennessee Office of Research and Engagement, and (4) the University of Tennessee Department of Mechanical, Aerospace and Biomedical Engineering start-up funds. The authors have no conflicts of interest to disclose.

